# Source-to-Target Automatic Rotating Estimation (STARE) – a publicly-available, blood-free quantification approach for PET tracers with irreversible kinetics: Theoretical framework and validation for [^18^F]FDG

**DOI:** 10.1101/2021.09.15.460504

**Authors:** Elizabeth A Bartlett, R Todd Ogden, J John Mann, Francesca Zanderigo

**Affiliations:** Molecular Imaging and Neuropathology Area, New York State Psychiatric Institute, New York, USA; Department of Psychiatry, Columbia University Medical Center, New York, USA; Department of Biostatistics, Mailman School of Public Health, Columbia University Medical Center, New York, USA; Department of Radiology, Columbia University Medical Center, New York, USA

**Keywords:** blood-free PET quantification, irreversible radiotracers, net influx rate, kinetic modeling, source-to-target modeling

## Abstract

**Introduction:** Full quantification of positron emission tomography (PET) data requires an input function. This generally means arterial blood sampling, which is invasive, labor-intensive and burdensome. There is no current, standardized method to fully quantify PET radiotracers with irreversible kinetics in the absence of blood data. Here, we present Source-to-Target Automatic Rotating Estimation (STARE), a novel, data-driven approach to quantify the net influx rate (K_i_) of irreversible PET radiotracers, that requires only individual-level PET data and no blood data. We validate STARE with [^18^F]FDG PET and assess its performance using simulations.

**Methods:** STARE builds upon a source-to-target tissue model, where the tracer time activity curves (TACs) in multiple “target” regions are expressed at once as a function of a “source” region, based on the two-tissue irreversible compartment model, and separates target region K_i_ from source K_i_ by fitting the source-to-target model across all target regions simultaneously. To ensure identifiability, data-driven, subject-specific anchoring is used in the STARE minimization, which takes advantage of the PET signal in a vasculature cluster in the FOV that is automatically extracted and partial volume-corrected. To avoid the need for any *a priori* determination of a single source region, each of the considered regions acts in turn as the source, and a final K_i_ is estimated in each region by averaging the estimates obtained in each source rotation.

**Results:** In a large dataset of [^18^F]FDG human scans (N=69), STARE K_i_ estimates were in good agreement with corresponding arterial blood-based estimates (regression slope=0.88, r=0.80), and were precisely estimated, as assessed by comparing STARE K_i_ estimates across several runs of the algorithm (coefficient of variation across runs=6.74 ± 2.48%). In simulations, STARE K_i_ estimates were largely robust to factors that influence the individualized anchoring used within its algorithm.

**Conclusion:** Through simulations and application to [^18^F]FDG PET data, feasibility is demonstrated for STARE blood-free, data-driven quantification of K_i_. Future work will include applying STARE to PET data obtained with a portable PET camera and to other irreversible radiotracers.

## 1 INTRODUCTION

Positron emission tomography (PET) allows for *in vivo* quantification of brain metabolism of different molecules and other neurotransmitter system components such as receptors, enzymes and ion channels. Full quantification of dynamically-acquired PET data can provide estimates of the amount of radiotracer that is specifically bound to a target of interest in the brain via binding potential options such as BP_F_ and BP_P_^1^ in the case of radiotracers with reversible kinetics (e.g., [^11^C]UCB-J and [^11^C]PBR28). We can quantify the total radiotracer uptake and metabolism via estimation of K_i_, the net influx rate of radiotracer into tissue from the vascular compartment, in the case of radiotracers with irreversible kinetics (e.g., [^18^F]FDG and [^18^F]FDOPA)^1^. Essential to obtaining these quantitative estimates, whether through kinetic compartment modeling or graphical approaches (e.g., Patlak plot^2^), is knowing the input function. This is the concentration of radiotracer and its radiometabolites in the blood compartment throughout the scan, which is used to inform the modeling^1^. The best validated and most widely used source of an input function is arterial blood. While arterial blood sampling via arterial catheterization has been safely applied in numerous research studies, it adds patient burden, cost and is labor intensive when employed in PET protocols. Therefore, efforts to develop, validate, and disseminate alternative, less-invasive quantification techniques for PET imaging would, if successful, enhance use of fully quantitative PET in both research and clinical settings. Here, we present Source-to-Target Automatic Rotating Estimation (STARE), a novel approach that performs full PET quantification of K_i_ for a PET radiotracer with irreversible kinetics, using only the individual-level PET brain data in a completely data-driven manner, without requiring collection of blood. We introduce here the theory behind this approach, and report its initial implementation and validation in 69 previously acquired [^18^F]FDG scans in humans^3,4^ and in [^18^F]FDG-based simulations.

[^18^F]FDG, a glucose analog, is the most ubiquitously used radiotracer in PET imaging and yields information on glucose metabolism, which in the brain is considered a marker for neural activity^5^. Due to the requirements for full quantification of [^18^F]FDG data to estimate K_i_ and the corresponding metabolic rate of glucose (CMR_glu_)^6,7^, semi-quantitative metrics, such as the standardized uptake value (SUV), have been frequently employed instead^8^. However, without strict standardization of SUVs^9–11^, quantification based on estimating K_i_ and CMR_glu_ is preferable, especially where highly sensitive PET metrics are required to detect subtle biological differences. Therefore, there has been much work to develop methods that can estimate K_i_ and CMR_glu_ without relying on an arterial input function (AIF). Common classes of less invasive or non-invasive quantification techniques include reference region approaches, image-derived input functions (IDIFs), population-based input functions (PBIFs), and simultaneous estimation (SIME) of the input function^12–21^.

Reference region approaches quantify PET outcome measures with respect to the tracer time activity curve (TAC) in a region assumed to be devoid of the target of interest for reversible or irreversible tracers^20^. However, for many tracers, including [^18^F]FDG, there is no such target-free brain region. For example, glucose is taken up by all living tissues, and this precludes application of reference region approaches.

Other proposed less-invasive methods, IDIFs, PBIFs and SIME, generally seek to recover a proxy for the AIF, where the AIF proxy is typically “anchored” or scaled for the individual in question, commonly by using one or more blood samples. Obviously, such an approach does not entirely eliminate the need for blood sampling during scanning.

IDIFs rely on extracting the radioactivity within vasculature within the PET field of view (FOV) and PBIFs rely on blood data previously acquired with the same tracer in other subjects, but both still currently require individual blood-based anchoring for practical application. SIME of the input function achieves less-invasive quantification by fitting the proper tissue compartment model to multiple brain regions’ TACs simultaneously. This allows the free parameters of the model, that is the parameters requiring estimation, to be estimated simultaneously, under the usual assumption that the AIF is the input function common to all regions^12–14,16,17,22–26^. In the case of SIME of the input function, the free parameters are those describing the kinetics of the tracer in the tissue regions (e.g., in the case of an irreversible tracer, the micro-parameters K_1_, k_2_, and k_3_ for each region) and those describing the AIF curve (e.g., the parameters in the model used to describe the arterial plasma curve, which is often described as the sum of three decreasing exponentials). However, to ensure identifiability of all free parameters at the individual level, SIME of the input function also requires “anchoring” the solution using at least one blood sample acquired during scanning^16^.

Although theoretically, one single arterial sample for IDIFs, PBIFs, or SIME could be acquired using an arterial puncture, the procedure can generate a sudden reaction in the subject under scanning, causing head motion and alterations in blood pressure and cerebral blood flow that may impact tracer delivery to and washout from the brain. Thus, it is preferable to avoid collecting even a single arterial sample. For [^18^F]FDG, where the radioactivity in venous blood approximates closely the radioactivity in arterial blood beyond 40 minutes post-injection^27,28^, SIME, IDIFs, and PBIFs have been applied using one or more venous plasma samples acquired late in the scan for anchoring^12,14,15,29–34^. This approach still requires placement of a second intravenous catheter, in addition to the line used for radiotracer injection, and then counting of venous blood activity in a well counter, adding complexity to PET acquisition.

There has been some success for completely noninvasive full quantification of PET data that obviates the need for any blood collection during the scan. These solutions often require combinations of multiple techniques to achieve acceptable performance. One solution anchors a PBIF with IDIF information derived with whole-body PET scanning^35^. Deep learning has also been leveraged to obtain blood-free quantification for [^18^F]FDG; however, to our knowledge this has yet to be validated with human scans^36^. Further, machine learning applied to precompiled electronic health record (EHR) data has been combined with SIME of the input function to quantify [^18^F]FDG without the use of any blood samples^4^. However, these solutions are situation specific i.e., with whole body scanning, with large sets of biological variables in the form of EHR data, or with large datasets acquired from many subjects with the same radiotracer for training and validation of the machine learning algorithm.

We now propose STARE (Source-to-Target Automatic Rotating Estimation), a new, blood-free, data-driven approach to quantification of PET tracers with irreversible kinetics that relies only on individual-level dynamic PET data. STARE utilizes a source-to-target tissue model, where the tracer radioactivity curve in a “target” region is expressed as a function of a “source” region, to eliminate the dependency of compartmental modeling on arterial blood. This source-to-target tissue model must be adapted to allow us to disentangle the parameters of interest for the target region from those of the source region. We do this by considering multiple target regions at once, as a function of the common source, and fitting the source-to-target model across all target regions and the source simultaneously. This approach allows STARE to separate K_i_ in target regions from K_i_ in the source region. Differently from SIME of the input function, STARE does not use data from blood samples to “anchor” the solution to the given individual but instead, uses bootstrapped, PET image-derived measures of concentration in the vasculature present in the FOV, as described below. We have validated STARE in a large set of human [^18^F]FDG scans in comparison to AIF-based estimation and using simulations.

## 2 MATERIALS AND METHODS

### 2.1 Theoretical framework

STARE is based on a reformulation of the standard two-tissue irreversible compartment model (2TCirr). The standard 2TCirr model expresses the concentration of radiotracer in a target region of interest (*C_T_*(*t*)) as a function of the concentration of radiotracer in the arterial plasma (*C_p_*(*t*)) as follows:

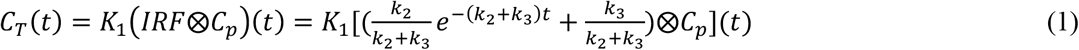

where *t* is time, *IRF* is the impulse response function for the target region, and *K*_1_, *k*_2_, *k*_3_ are the micro-parameters of the 2TCirr model for the target region. If *C_p_*(*t*) were available, fitting of the model in (1) to the TAC in a target region would result in estimates of the micro-parameters *K*_1_, *k*_2_, and *k*_3_, and thus, of *K_i_*, as *K_i_*, = *K*_1_*k*_3_/(*k*_2_ + *k*_3_).

Without acquisition of arterial plasma samples throughout the scan, an estimate of *C_p_*(*t*) is not available. In the case of a PET tracer with irreversible kinetics, such as [^18^F]FDG, Equation (1) typically holds for TACs from any brain region. Therefore, it is possible to reformulate Equation (1) so that the *C_T_*(*t*) in a target region is expressed as a function of the TAC in another region (*C_s_*(*t*)), which we denote here as “source”, thus obviating the need to know *C_p_*(*t*). This is similar to the substitutions performed in the case of reference region approaches for tracers with reversible kinetics, where in those cases, theoretical assumptions are made about the reference region: that it is devoid of specific binding to the target of interest. Differently from reference region approaches, however, in STARE, the only required theoretical assumption for the source region TAC is that it follows the 2TCirr model, as shown in Equations 2 and 3 below, where the source and target region TACs are both assumed to follow a 2TCirr model. As described in detail below, differently from reference region approaches, STARE’s simultaneous estimation of the parameters for multiple target regions at once allows estimation of “absolute” K_i_, estimates, rather than the “relative” outcome measures (BP_ND_) derived from reference region approaches.

By applying to Equations 2, shown below, Laplace transformation, substitution, and subsequent transformation back into the time-domain, we express each of the target region TACs as a function of its own micro-parameters, the micro-parameters describing the source region, and the source region TAC itself, as shown in Equation 3. Specifically, the TACs in both the target and source regions can be expressed in terms of the standard 2TCirr model as:

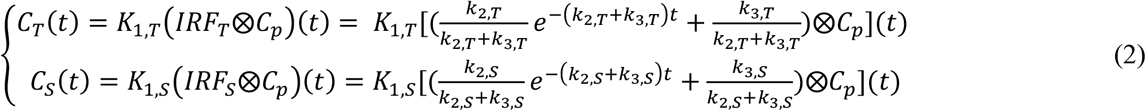

By applying the Laplace transform in the system of equations in (2) (see Appendix for full derivation), *C_p_*(*t*) can then be substituted out so that the target region TAC is modeled as:

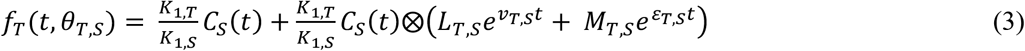

Which, we denote the source-to-target tissue model: where *t* is time, ⊗ denotes convolution, and *θ_T,s_* is the total set of free parameters to be estimated in STARE via optimization of the cost function in Equation 5. These free parameters are the 2TCirr micro-parameters (*K*_1_, *k*_2_, and *k*_3_) for each of the target regions and for the source region. As shown in Equation 3, *θ_T,s_* comprises the macro-parameters *L_T,s_*, *M_T,s_*, *v_T,s_*, and *ε_T,s_*, which are combinations of the 2TCirr free micro-parameters (see Appendix for full derivation).

However, applying the reformulation in Equation (3) for a single target and source region combination, only allows for quantification of PET outcome measures “relative” to the selected source region and thus, cannot yield absolute estimation of the target micro-parameters and of *K_i_*. This is analogous to reference region approaches estimating the relative outcome measure BP_ND_ in the case of radiotracers with reversible kinetics. In order to separate target *K_i_* from source *K_i_* to allow for absolute quantification, we follow a strategy analogous to simultaneous estimation (SIME) of the input function. In that context, the parameters describing the unknown AIF may be estimated together with the free parameters describing the tracer kinetics in the tissue by fitting all regions’ TACs at once, under the assumption that the AIF parameters are in common to all regions. Similarly, here we model multiple target regions at once, under the assumption that the source region parameters that they are expressed as a function of, are in common to all regions. The parameters for the target regions and the source region are then estimated at once with simultaneous estimation as follows.

Once each target region is expressed as a function of the source region (according to Equation (3)), the weighted sum of squared residuals is used across *N* target regions, where the residuals are the distances between each measured target TAC (*C_T_*(*t*)) (with *T* = 1,…,*N* indicating the different target regions) and the corresponding modeled target TAC (*f_T_*(*t*, *θ_T,s_*)) at each *t_m_* time point (*m* = 1,…,*n*), as follows:

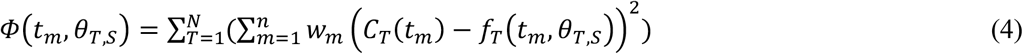

*w* in Equation (4) indicates a set of known weights for the different PET frame durations (as is standard in PET imaging).

Similarly to the case of SIME of the input function, however, minimization of Equation (4) will not yield unique estimates of the free parameters for both the target regions and source region. This is due to the fact that there exist multiple combinations of such free parameters that yield equivalently good TAC fits. In the case of SIME of the input function, this identifiability problem is solved by “anchoring” the solution to a blood sample acquired from the subject during scanning^13,16,37^.

Analogously, in STARE we ensure identifiability by effectively “anchoring” the estimation process, not to data from blood samples, but to PET-derived measures of activity in the vasculature present in the FOV (as are common with IDIF methods), as described in detail in the Implementation section. To do this, an additional penalty term is added to the weighted sum of squared residuals in Equation (4):

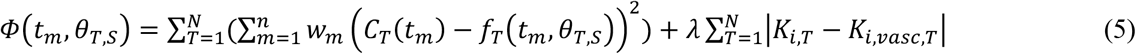

This term enforces identifiability by constraining the solution, and *K_i,T_*, the *K_i_* in region *T*, to a subject-specific neighborhood around estimates derived from the signal in the vasculature within the PET FOV (the *K_i,vasc,T_* values). As is described in 2.2.1 *Implementation: STARE Anchoring*, this is estimated in a data-driven manner based on the individual’s PET data. The PET signal is automatically extracted from vasculature within the PET FOV, as in IDIF techniques. However, the variability in kinetics across voxels within this vasculature region is then bootstrapped to generate a wide range of possible parameter estimates for each brain region for the given participant, from which the *K_i,vasc,T_* values are derived. The parameter *λ* is introduced to balance the contribution of the fit term and the penalty term in Equation (5), which may be advantageous for application to specific radiotracers. In this implementation *λ* is set to 1, as described in 2.2.2 *Implementation*.

Equation (5) requires designation of a “source” region. Although, any region whose TAC follows the 2TCirr model could be used as the source region, to ensure that final estimates do not rely on an arbitrary choice of source, we elected to allow each region to act in turn as the source region. Final *K_i_* estimates thus result from averaging the *K_i_* estimates obtained for each source “rotation”. The theoretical framework and implementation of STARE anchoring is described in Figure 1.

**Figure 1:**
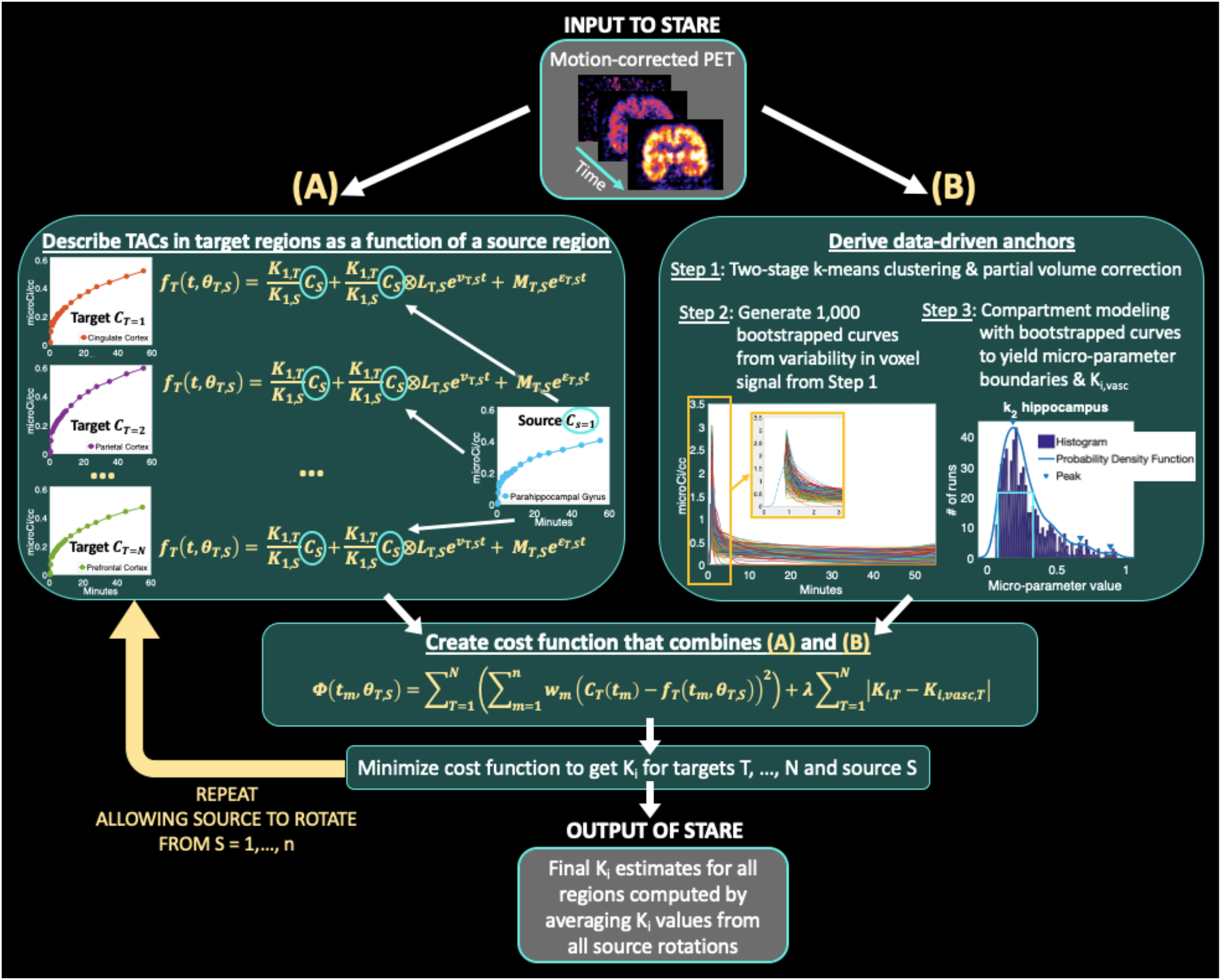
Graphical representation showing the theoretical framework and implementation of STARE

### 2.2 Implementation

STARE was applied to 69 [^18^F]FDG human brain scans^3,4,29^ and to simulated data considering TACs from six bilateral regions of interest (ROIs): cerebellum, cingulate cortex, hippocampus, parietal cortex, medial prefrontal cortex, and parahippocampal gyrus. STARE implementation is shown in Figure 1. Matlab 2016b (The MathWorks, Natick, MA) was used for implementation and all subsequent processing.

#### 2.2.1 STARE Anchoring

STARE anchors the estimation process to a unique solution for each individual via the additional penalty term in (5). Furthermore, data-driven boundaries are also generated to constrain the search space for all free parameters in the model. Here we describe how this anchoring can be fully automated.

We first use a two-step k-means clustering approach to automatically extract a vasculature cluster by parsing the dynamic PET data into characteristic regions, such as background, brain tissue with irreversible uptake, and vasculature, in a completely data-driven manner. This is followed by partial volume correction (PVC) of the final vasculature cluster (Figure 1). To anchor the STARE solution in the correct subject-specific “neighborhood” of the free parameters space, we capitalize on the variability of signal within the extracted vasculature cluster, rather than extracting a single summary metric, using bootstrapping of the voxel TACs within the vasculature cluster. More specifically:

##### 2.2.1.1 K-means Step 1

To automatically select the optimal number of clusters to be extracted, based solely on an individual’s PET data, rather than *a priori* assuming that a pre-set number of clusters will optimally partition all scans, k-means clustering runs multiple times, each with a different number of extracted clusters. For [^18^F]FDG, we used a generous range from 6 to 40 clusters. An optimal vasculature cluster is then automatically selected from each k-means runs by: (1) eliminating all clusters where the maximum value of the average TAC within the cluster corresponds to the end-point of the curve because that indicates an irreversible kinetic, which more likely represents tissue; (2) eliminating all clusters whose average TAC shows negative values because that most likely represents background voxels; and (3) selecting from the remaining clusters, the one whose average TAC shows the highest peak value at the earliest time of peak because that most likely represents PET signal arising from blood vasculature.

##### 2.2.1.2 K-means Step 2

K-means clustering then runs again only on those voxels within the cluster selected during Step 1, with the assumption that this cluster represents a gross estimate of the vasculature within the FOV, which might be corrupted by some spill-in from nearby tissue (especially late in the scan) and spill-out of vasculature signal (especially early in the scan). Because it might be that this initial gross estimate is comprised of signal from arteries, veins, sinuses, and tissue, in our implementation, we elected to extract 4 clusters during Step 2. Among the extracted clusters, similar to Step 1, the voxels belonging to clusters whose average TAC has the highest peak value are selected as the final vasculature cluster. Prior studies have shown that with [^18^F]FDG, as well as with a multitude of other radiotracers and drugs, the signal arising from the arterial vasculature early in the scan is higher than the signal arising from the venous vasculature, as well as any other tissues^29,37–39^. Therefore, the k-means cluster with the highest early-scan peak is likely to be the closest approximation to the true activity in arterial blood.

##### 2.2.1.3 Partial Volume Correction

PVC via Single Target Correction (STC) is then applied to voxels within the final vasculature cluster from Step 2^40^. STC was previously optimized and validated, where PVC is performed on a voxel-wise basis for a single region (i.e., the final vasculature cluster), accounting for the voxel-wise spill-in of radioactivity into the vasculature cluster and spill-out of radioactivity from the vasculature cluster^40^. Given the reconstruction parameters of the [^18^F]FDG dataset considered here, a point-spread function of 5.9 mm full width at half maximum (FWHM) was used for PVC via STC^3,40^. This parameter should be selected and optimized based on the scanner resolution and reconstruction parameters (e.g., postreconstruction smoothing) for the dataset at hand.

##### 2.2.1.4 Data-driven extraction of Equation (5) penalty term and parameter space bounds

The partial volume corrected TACs from each voxel within the final vasculature cluster are used to derive an individualized, data-driven penalty term (see Equation (5)), and to derive bounds for the model free parameters. As described in detail below, we leveraged the variability of the TACs within the partial volume-corrected final vasculature cluster to extract *K_i,vasc_* values for each target region *T*. Our new approach for estimating *K_i,vasc,T_* extends beyond standard IDIF approaches that often simply use the average TAC in a vasculature cluster, because these standard approaches typically require blood-based scaling^41^.

First, we simulate many instances of “IDIF” curves (*C_IDIF,b_*(*t*), with *b* = 1,…,*B* and *B* = 1000 in our implementation), by starting from the average TAC in the vasculature cluster, and bootstrapping curves that fall between one standard deviation below and one above the average TAC, according to the following formula:

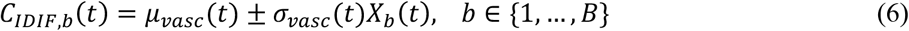

where *t* is time, *μ_vasc_*(*t*) is the average TAC across the voxels in the vasculature cluster, *σ_vasc_*(*t*) is the standard deviation across the voxel TACs in the vasculature cluster, and *X_b_* is a uniformly distributed random number in the interval [0,1] at each time *t* for each bootstrapping iteration *b*. Each of the generated “IDIF” curves is then fitted with a 3-decreasing exponential model (*F_IDIF,b_*(*t*)), commonly adopted in PET to describe the post-peak blood input function, and held at *μ_vasc_*(*t*) from time zero to the time of peak, to avoid any non-physiological uptake patterns at the very beginning of the scan (see Figure 1 for an example of the generated bootstrapped curves).

Each *F_IDIF,b_*(*t*) curve then serves as a proxy for the input function, allowing each of the regions’ TACs to be fit with the 2TCirr model. This approach yields *B* sets of estimated 2TCirr micro-parameters per region, from whose kernel, density estimate can be obtained via nonparametric estimation of the probability density function (Matlab function “ksdensity”). See Figure 1 for an example of the *k*_2_ density estimate in hippocampus. For each micro-parameter (*K*_1_, *k*_2_, and *k*_3_) in each region, the bounds constraining the search space during optimization of Equation (5) ([*LB*, *UB*]) are then automatically set using the FWHM of the estimated probability density estimate (*Pr_k_*), such that in each region:

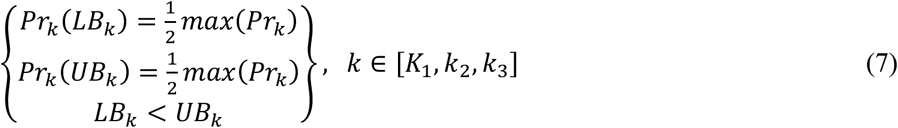

See Figure 1 for an example of [*LB*, *UB*] for hippocampal *k*_2_ values.

The *B* sets of estimated 2TCirr micro-parameters are then also used to compute *B* corresponding *K_i_* values, from whose distribution, probability density function is also derived. The *K_i,vasc_* value used to penalize Equation (5), is then derived for each region as 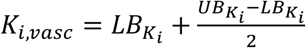.

#### 2.2.2 STARE Cost Function Minimization

For each source rotation, Equation (5) is minimized in parallel via simulated annealing^42^. Simulated annealing was implemented using the Matlab function “simulannealbnd” with default options for initial temperature (100), reannealing interval (100), function tolerance (1e-6), and maximum iterations (Inf). The simulated annealing initial guesses were set randomly in the range [*LB*, *UB*] for each parameter, and the search space confined to [*LB*, *UB*] for each parameter.

In our [^18^F]FDG data, we find that the magnitude of the distance between measured and modeled TACs (fit term) in Equation (5) is comparable to the magnitude of the difference between *K_i_* estimates (penalty term) and thus, in this implementation, we set *λ* to 1.

#### 2.2.3 Vascular Correction

The source-to-target tissue model in Equation (3) assumes that the TACs of target and source regions are corrected for vascular contribution. However, in the absence of measurements of the radiotracer activity in whole blood or plasma, such vascular correction is not easily achieved. We, therefore, implemented an optional vascular correction procedure within STARE. The partial volume corrected average TAC in the vasculature cluster (*μ_vasc_*(*t*)) can be used to perform vascular correction of the TACs in all regions according to the following:

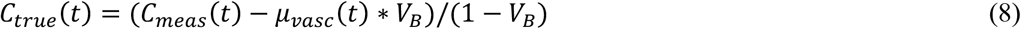

where *t* is time, *C_meas_*(*t*) is the measured tissue radioactivity curve from the PET camera, *C_true_*(*t*) is the tissue radioactivity corrected for vascular contribution, and *V_B_* is a user-modifiable vascular volume fraction. We investigated the effect of neglecting vascular correction (*V_B_* = 0.00), and including STARE’s implementation of vascular correction (*V_B_* = 0.05^43^), according to Equation (8), on STARE performance in quantifying *K_i_*.

### 2.3 AIF-based quantification of the available [^18^F]FDG dataset

An available set of 69 previously acquired and published [^18^F]FDG scans^3,4,29^ was considered, which included participants with mild cognitive impairment, mild Alzheimer’s disease and healthy controls. Per the data sharing agreement, data could be made available by request to Drs. J John Mann/Davangere P Devanand. As previously described, written informed consent was obtained from all participants^3,4,29^ and the study was approved by the Institutional Review Board of the New York State Psychiatric Institute and Columbia University. Acquisition details are previously described^3^. Arterial plasma was sampled throughout the scan via arterial catheterization as previously described^44^. To generate a “gold-standard” AIF, the measured values of tracer total radioactivity in arterial plasma were interpolated from time 0 to the time of the plasma peak. After the peak, a sum of three decreasing exponentials was used to fit the radioactivity data via non-linear least squares. The AIF then was used as the input function to the 2TCirr compartmental model to fit each considered TAC, estimate the model micro-parameters, and calculate corresponding *K_i_* estimates. These blood-based 2TCirr *K_i_* estimates were considered to be the “gold-standard” comparison for STARE-based estimates of *K_i_*, although we acknowledge that AIF data, and thus AIF-based *K_i_* estimates, may be prone to noise and error^45^. For comparison with STARE, AIF-based 2TCirr was run either while neglecting vascular correction (*V_B_* = 0.00) or with standard vascular correction, using the plasma AIF (because the radioactivity curve in whole blood was not measured in this previously acquired dataset) with *V_B_* = 0.05^43^, according to Equation (8).

### 2.4 Assessing STARE performance relative to AIF-based quantification

#### 2.4.1 STARE Accuracy

Agreement between STARE *K_i_* and AIF-based 2TCirr *K_i_* estimates was assessed using linear regression and Pearson correlation. Regressions and correlations were computed for: (1) all regions and scans together, (2) scan by scan, across all regions, and (3) region by region, across all scans. Signed percent difference was also computed as: (STARE *K_i_* – 2TCirr *K_i_*)/2TCirr *K_i_*\*100. To assess the possibility of regional dependence in STARE performance, a linear mixed effects model was fit with outcome = the natural logarithm of *K_i_* and fixed effects = region and quantification method (STARE vs. AIF-based 2TCirr). Participant was modeled as a random effect.

#### 2.4.2 STARE Precision

Because STARE contains non-deterministic algorithms (i.e., k-means clustering and simulated annealing), the stability in estimating *K_i_* was tested. STARE was run 10 times per scan for a random subset of the N=69 [^18^F]FDG scans. Stability of STARE across runs and regions was assessed with the coefficient of variation of *K_i_* estimates (COV = standard deviation / mean). All statistics were performed in R version 4.0.3^46,47^.

#### 2.4.3 STARE Performance Across Diagnostic Groups

Given that the [^18^F]FDG dataset includes a transdiagnostic sample, we assessed whether STARE’s performance varies with disease states (participants with mild cognitive impairment (MCI), mild Alzheimer’s disease (AD) and healthy controls (HC)). A linear mixed model was fit as in 2.4.1, but with diagnostic group added as a fixed factor (in addition to region and quantification method). The two-way interaction of diagnosis by method to test if K_i_ varied by quantification method on a diagnosis-specific basis was examined.

### 2.5 Simulations

A set of simulation studies was designed to examine the sensitivity of STARE to variations in the procedure used to determine *K_i,vasc,T_* and the upper and lower bounds ([*LB*, *UB*]) for the model free parameters. One set of simulations (A) investigated the effect of variations in *μ_vasc_*(*t*) (the average TAC across the voxels in the final vasculature cluster) by manipulating its area under the curve (AUC), the curvature of its tail, and its overall shape (in this last case, while holding the AUC constant). Another set of simulations (B) investigated the effect of variations in *σ_vasc_*(*t*) (the time-wise standard deviation of voxel activities within the final vasculature cluster). Across all simulations, the same representative [^18^F]FDG scan was used as a starting point. To quantify how much each simulation altered *μ_vasc_*(*t*) or *σ_vasc_*(*t*) relative to the original curves, a difference score was computed as the absolute summed difference across time-points between the original *μ_vasc_*(*t*) or *σ_vasc_*(*t*) and the manipulated *μ_vasc_*(*t*) or *σ_vasc_*(*t*) curves. This was compared to the percent difference in *K_i_* between the original STARE run and the STARE run following the given manipulation.

#### 2.5.1 Effects of *u_vasc_*(*t*) on STARE performance

<u>2.5.1.1:</u> The AUC of *μ_vasc_*(*t*) was manipulated, while maintaining the shape of the curve, by solving for *add_var_* in the following equation:

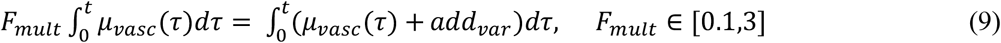

where *t* is time and *F_mult_* is the scaling factor that alters the AUC (with *F_mult_* in the range [0.1, 3]). *add_var_* was estimated at each *F_mult_* instance, and then added to the representative scan’s *μ_vasc_*(*t*).

<u>2.5.1.2:</u> The curvature in the tail of *μ_vasc_*(*t*) was manipulated by altering the exponential decay of the tail. To do this, the extracted *μ_vasc_*(*t*) was fitted to the usual 3 decreasing exponential model, and the exponential term with the smallest decay constant, which models the late-scan kinetics of *μ_vasc_*(*t*), was modulated using a multiplicative scaling factor to increase or decrease the rate of decay. The same *F_mult_* values as in A.1 were used. The corresponding AUC of the simulated curve was allowed to change with the different *F_mult_* values.

<u>2.5.1.3:</u> The shape of the *μ_vasc_*(*t*) curve was then manipulated, while holding the AUC constant, by using the same *μ_vasc_*(*t*) fit as in A.2. Sets of three decay constants, one per each decreasing exponential, were randomly generated in each simulation and combined with the original corresponding initial magnitude values from A.2. This new simulated curve was then divided by a factor, *div_var_*, that was estimated in each iteration using a similar procedure to Equation (9), in order to hold the final simulated curve’s AUC constant at the representative scan’s original value.

#### 2.5.2 Effects of *σ_vasc_*(*t*) on STARE performance

The standard deviation of the tracer radioactivity in the voxels within the final vasculature cluster, *σ_vasc_*(*t*), was then manipulated via additive scaling using the same approach as in A. 1 (Equation (9)). In this case, however, *σ_vasc_*(*t*) substituted for *μ_vasc_*(*t*) in Equation (9) and the estimated *add_var_* at each *F_mult_* was added to the representative scan’s *σ_vasc_*(*t*).

## 3 RESULTS

### 3.1 STARE Accuracy

Blood-free STARE *K_i_* estimates were highly correlated with AIF-based *K_i_* estimates (regression slope (b)=0.88, y-intercept=0.004, Pearson’s r=0.80, *p*<0.001, Figure 2(A), Table 1)). Although the regression slope was less than one, the intercept was greater than zero and STARE K_i_ estimates were on average modestly overestimated relative to AIF-based estimates (signed percent difference: 5.07% ± 18.14%; Table 1). STARE K_i_ estimates were 0.00091 ± 0.0041 greater than AIF K_i_ (Figure 2(B)).

**Figure 2:**
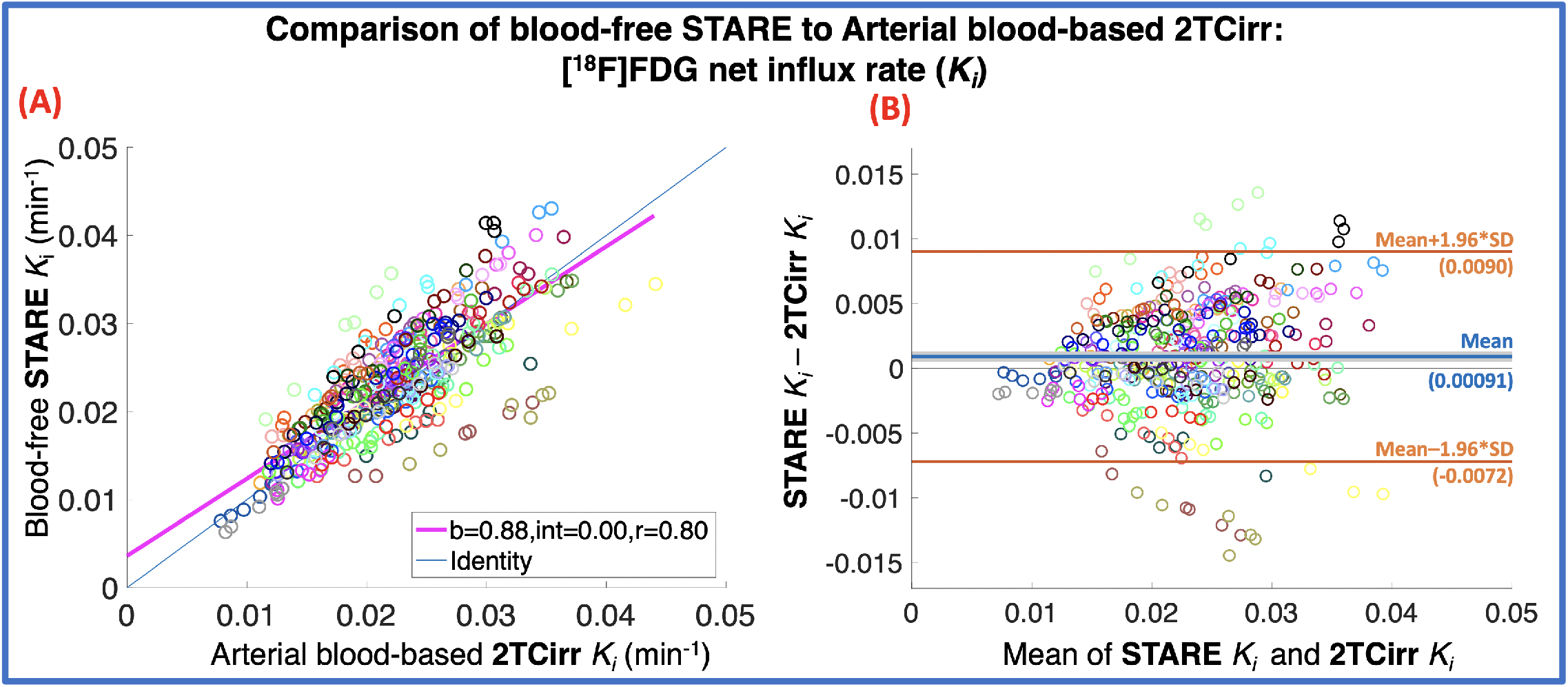
Blood-free STARE K_i_ vs. K_i_ obtained via arterial blood-based two-tissue irreversible (2TCirr) compartmental modeling in 69 [^18^F]FDG scans. In both (A) & (B): Each color corresponds to a single scan, where K_i_ is quantified for 6 regions (cerebellum, cingulate cortex, hippocampus, parahippocampal gyrus, parietal cortex, and prefrontal cortex). (A) Scatter plot, with linear regression slope, intercept, and Pearson’s correlation coefficient reported across all subjects and regions. (B) Bland-Altman plot with mean K_i_ difference (i.e., overall bias) shown in blue and the 95% confidence interval for the mean difference estimate shown in light grey; limits of agreement (95% confidence intervals of mean difference) are shown in orange.

**Table 1:** STARE Ki Performance Compared to 2TCirr *K_i_*.

| Table 1: STARE $K_i$ Performance Compared to 2TCirr $K_i$ | | | | |
| --- | --- | --- | --- | --- |
| | Regression Slope (b) | Intercept | Pearson's r | Percent Difference<br>(STARE $K_i$ – 2TCirr $K_i$) / 2TCirr $K_i$ * 100<br>(mean ± standard deviation) |
| Across all regions<br>(Figure 2(A)) | 0.88 | 0.00 | 0.80 | $5.07 \pm 18.14$ |
| Cerebellum | 0.71 | 0.01 | 0.66 | $4.83 \pm 18.3\%$ |
| Cingulate cortex | 0.80 | 0.01 | 0.68 | $4.83 \pm 18.0\%$ |
| Hippocampus | 0.83 | 0.00 | 0.71 | $5.97 \pm 18.7\%$ |
| Parietal cortex | 0.82 | 0.01 | 0.72 | $4.63 \pm 17.83\%$ |
| Medial prefrontal cortex | 0.85 | 0.00 | 0.72 | $4.82 \pm 18.18\%$ |
| Parahippocampal gyrus | 0.72 | 0.01 | 0.66 | $5.36 \pm 18.44\%$ |

Within individual scans, across the six regions considered, the agreement between STARE and AIF-based *K_i_* estimates was assessed via regression, and slopes ranged from b=0.59 to 1.58, while correlation coefficients ranged from r=0.95 to 1.00 (all *p*<0.005). Although the Pearson’s r values ranged from 0.66 to 0.72 and the slopes ranged from 0.71 to 0.85 for the comparison of individual regions’ STARE K_i_ estimates vs. AIF-based K_i_ estimates region-by-region, across all subjects (Table 1), there was no significant statistical evidence for a regional dependence in STARE’s estimation of K_i_ relative to AIF-based K_i_ estimates (p=0.999).

### 3.2 STARE Precision

Although STARE includes non-deterministic algorithms, STARE stably estimated *K_i_* across runs (COV = 6.74 ± 2.48%).

### 3.3 STARE Performance Across Diagnostic Groups

We tested whether STARE’s estimation of K_i_ was consistent across diagnostic groups (AD, MCI, and HC). The interaction between diagnosis (AD vs MCI vs HC) and quantification method (STARE vs AIF-based) was non-significant (p=0.17), indicating that the difference in K_i_ values estimated with the two methods did not significantly vary across diagnoses.

### 3.4 STARE Vascular Correction

While the level of correlation between STARE K_i_ and corresponding AIF-based estimates is not affected by whether the vascular correction strategy is considered (r=0.78) or not within STARE (r=0.79), their agreement slightly varies. More specifically, when comparing STARE K_i_ estimates obtained without the vascular correction strategy to AIF-based K_i_ estimates obtained with V_B_ = 5%, the slope of the regression line is b=0.83, (Figure 3(A)). When the proposed vascular correction strategy is used within STARE and the same comparison is made, the slope of the regression line is b=0.87 (Figure 3(B)), suggesting that considering the vascular correction strategy is favorable, at least in this [^18^F]FDG dataset.

**Figure 3:**
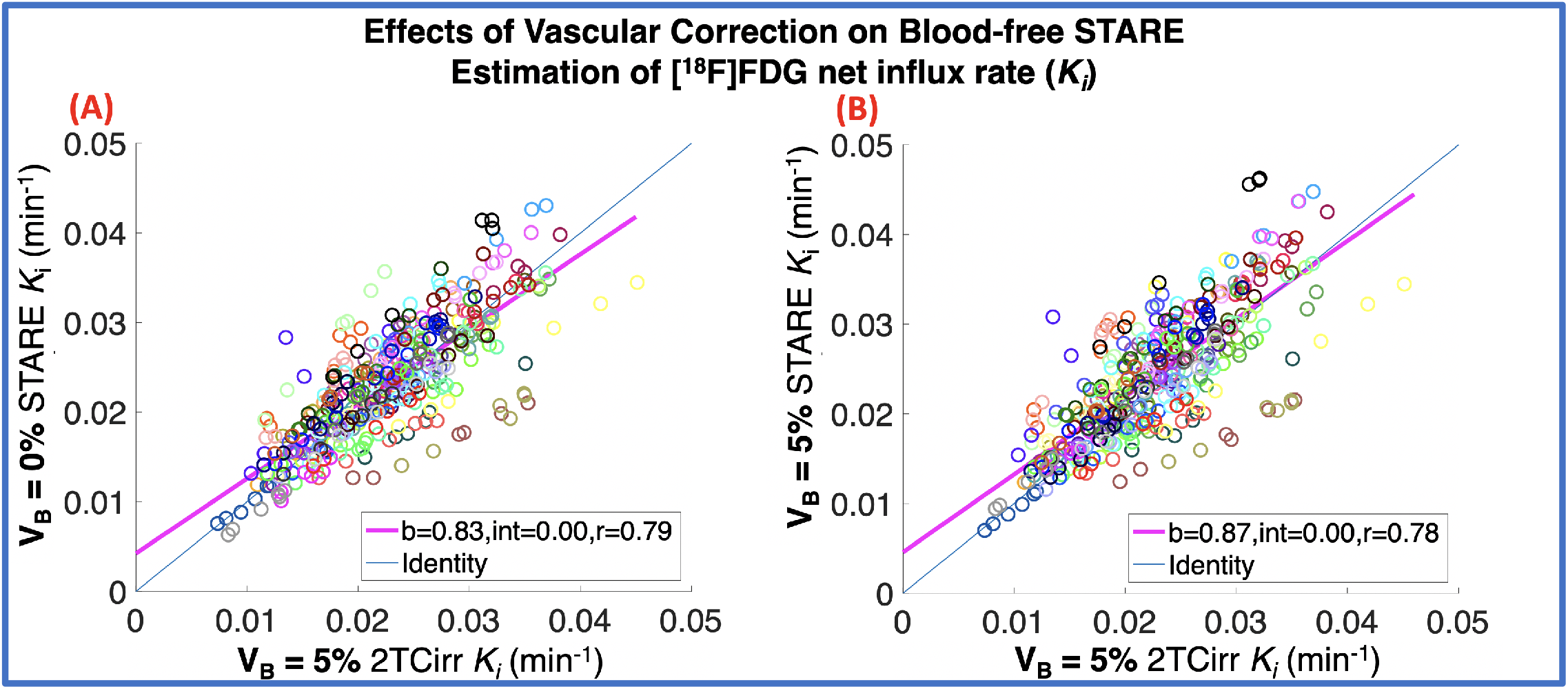
Effect of Vascular Correction on STARE estimates of K_i_. In both (A) & (B): Each color corresponds to a single scan, where K_i_ is quantified for six regions (cerebellum, cingulate cortex, hippocampus, parahippocampal gyrus, parietal cortex, and prefrontal cortex). Linear regression results and Pearson’s correlation coefficients are shown. (A) STARE K_i_ estimates with V_B_ = 0% (no vascular correction, y-axis) relative to AIF-based 2TCirr K_i_ estimates with V_B_ = 5% (x-axis) and (B) STARE K_i_ estimates with V_B_ = 5% (within STARE vascular correction, y-axis) relative to AIF-based 2TCirr K_i_ estimates with V_B_ = 5% (x-axis).

### 3.4 STARE Simulations

Simulation showed that altering the overall amplitude of signal within the vasculature cluster used to generate the STARE anchors (i.e. altering the AUC of *μ_vasc_*(*t*) via additive scaling) had the most substantial impact on STARE *K_i_* estimation (*Simulation A.1*; Figure 4.A. 1). To compare with human data, in the 69 [^18^F]FDG scans, the mean percent difference in AUC between STARE *μ_vasc_*(*t*) and the AIF was −6.10 ± 15.69%. According to our simulations, when the simulated difference in the AUC of *μ_vasc_*(*t*) was 10%, it yielded a −9.97% change in *K_i_* estimates.

**Figure 4:**
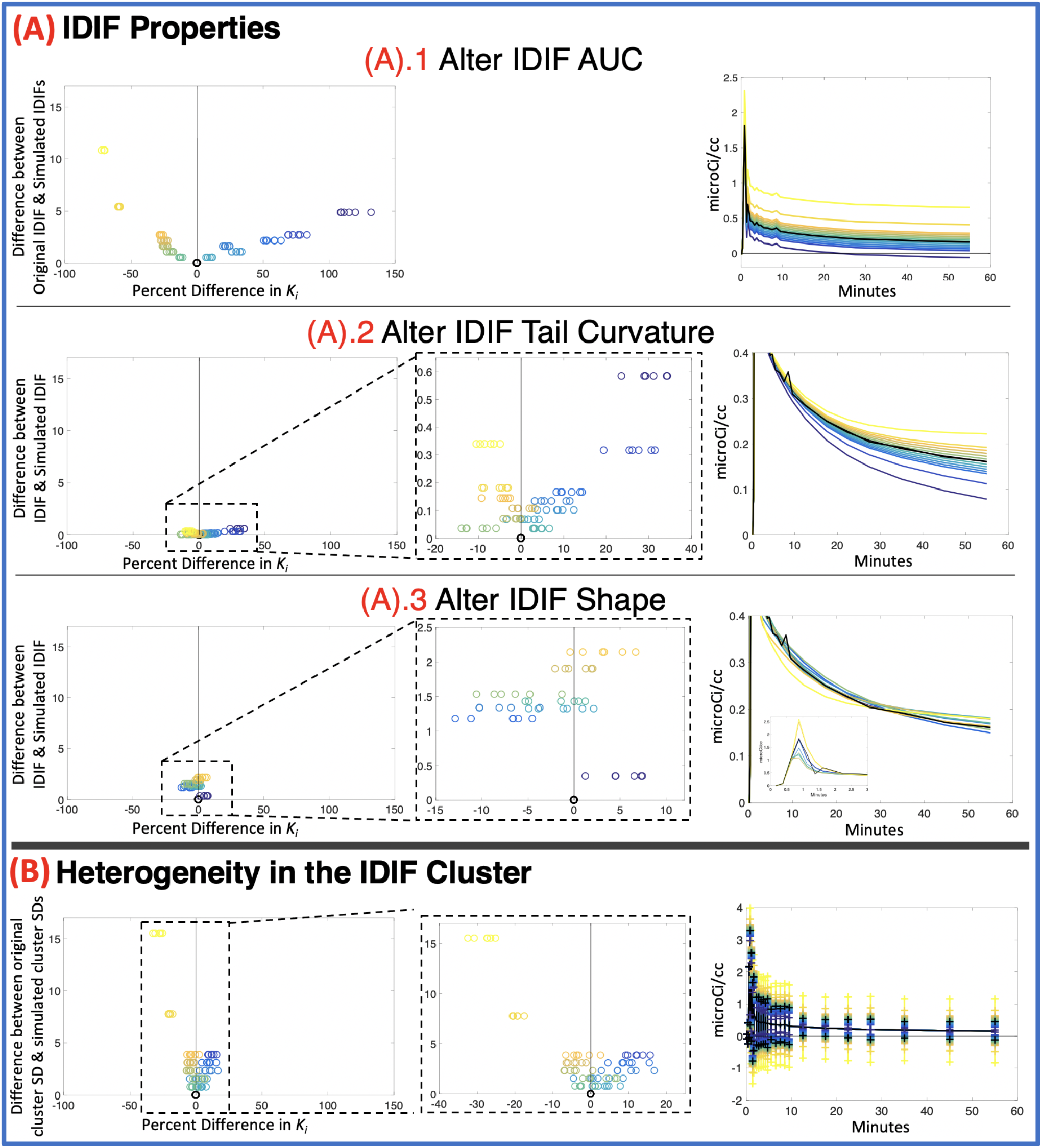
Simulation results for STARE. The data from a single, representative scan were used to build all simulations in (A) and (B). Simulations A.1 through A. 3 alter scaling and shape characteristics of the signal in the vasculature cluster (i.e. the mean of voxel radioactivities in the vasculature cluster, μ_vasc_(t)), whereas Simulation B alters the standard deviation of the radioactivity of voxels within the vasculature cluster (σ_vasc_ (t)). These properties are used in the generation of the STARE anchors. The left columns of (A) and (B) use the same metric on the y-axes to assess the difference between the original and simulated μ_vasc_(t) or σ_vasc_(t) curves, which sums the difference between the curves across all time-points. The x-axes of the left columns of (A) and (B) are the percent difference in STARE-estimated K_i_ between that simulation iteration and the original result. If necessary, a center column is shown as a zoomed-in inset of the left column. The right columns of (A) and (B) show the simulated curves (with the original curves shown in black)). Within each simulation (across rows), the colors (on a yellow-to-blue scale) are consistent from left to right.

STARE was relatively robust to alterations in the kinetics of signal arising from the vasculature cluster (*Simulation A. 2 and A. 3*; Figure 4.A.2 and 4.A.3) and the level of voxel-wise variance in the vasculature cluster used to generate the STARE anchors (*Simulation B*; Figure 4.B).

More specifically across all simulations, *K_i_* only changed by greater than 50% with respect to the original, non-simulated STARE *K_i_* values in *Simulation A.1: μ_vasc_*(*t*) *AUC* under the following simulated conditions: (1) *μ_vasc_*(*t*) AUC was increased by at least 200% (corresponding to a y-axis difference score of at least 5.40 in Figure 4.A.1) or (2) *μ_vasc_*(*t*) AUC was reduced by at least 40% (corresponding to a y-axis difference score of at least 2.16 in Figure 4.A.1). From Figure 4.A.1, we can observe that: (1) altering *μ_vasc_*(*t*) AUC yielded the expected inverse effects on STARE *K_i_* estimates, such that, for example, increasing the AUC yielded negative biases in STARE *K_i_* (Figure 4.A.1); and (2) reducing *μ_vasc_*(*t*) AUC had a greater impact on STARE *K,* than increasing *μ_vasc_*(*t*) AUC, presumably due to instabilities in the 2TCirr modeling in the free parameter bound generation because the input function tail approached 0 microCi/cc in activity and at times became negative.

In *Simulation A.2*, which manipulated the kinetics of the amplitude of signal arising from the vasculature cluster (allowing the AUC to change with change in shape), the simulated *K_i_* estimates changed by less than 50%, even when the rate of IDIF tail decay was increased by 300% or decreased by 90% (Figure 4.A.2). In *Simulation A*. 3, where the AUC of *μ_vasc_*(*t*) was held constant, while the shape was varied, a much smaller impact on STARE *K_i_* was observed, with all percent differences less than 15% (Figure 4.A.3). Similarly, altering the level of voxel-wise variance within the vasculature cluster (*Simulation B*), had little impact on STARE *K_i_* (Figure 4.B).

## 4 DISCUSSION

In this study, we present the theory and an initial validation with human [^18^F]FDG scans, for a novel, publicly available PET quantification approach – STARE – that quantifies the net influx rate (K_i_) of radiotracers with irreversible kinetics (STARE will be available on GITHUB and link will be provided here pending paper acceptance). This method, in theory, allows for noninvasive PET quantification that does not rely on blood sampling during the course of a PET scan and operates in an automatic, data-driven, individual-subject manner. To summarize our findings, in a large dataset of [^18^F]FDG scans in humans^3^: (1) K_i_ values obtained via STARE showed, on average, modest overestimation and were strongly correlated with “gold-standard”, AIF-based K_i_ estimates, and (2) K_i_ values obtained via STARE were stably and thus, precisely estimated. In simulations, K_i_ was largely robust to deviations in STARE anchoring.

The goal of STARE is to provide blood-free full quantification of data acquired with PET radiotracers with irreversible kinetics. While validated here using [^18^F]FDG, STARE’s theoretical framework is based on a manipulation of the general 2TCirr model. Therefore, theoretically, STARE can potentially be applied to any radiotracer whose kinetics can be fitted with an irreversible compartment model. Key considerations when validating STARE for another tracer include the heterogeneity of the PET signal across the brain and the level of noise present in the image according to the PET camera used to acquire the data. This validation, in comparison to AIF-based quantification, can be achieved with a modest sample size.

This highlights one of the advantages of STARE, which is that the approach does not depend on machine learning, for which large sample sizes are often required for adequate training of the algorithm. In fields like dynamic PET imaging, where data sharing initiatives are still in their infancy, and PET acquisition with arterial blood sampling is costly and complicated, large datasets meeting the appropriate requirements for machine learning are rare, especially for novel radiotracers. One such machine learning method, noninvasive SIME (nSIME), which was previously validated with the same [^18^F]FDG dataset considered here, trained a model with 83 different predictors extracted from precompiled EHR data to estimate [^18^F]FDG *K_i_* in conjunction with simultaneous estimation^4^. nSIME performed comparably well with blood-free STARE (r=0.80 STARE, r=0.83 nSIME^4^; Bland-Altman plots appear qualitatively comparable across methods, but nSIME appears to exhibit more bias), highlighting STARE performance even though it only requires an individual participant’s dynamic PET data without model training that is based on other participants’ data.

STARE performance was comparable across brain regions for both accuracy and precision. This finding highlights another key feature of STARE – the source rotation. Unlike reference region methods, where *a priori* knowledge is required to determine one appropriate region assumed to be devoid of radiotracer specifically bound to the target, STARE’s source rotation step does not require any *a priori* determination of a source region. Our findings show that STARE estimates K_i_ with equivalent accuracy when we compute final K_i_ estimates as averages across all rotations. We also provide preliminary evidence that STARE is robust to disease-specific uptake patterns, where we found that STARE performance did not significantly vary across HC, AD, and MCI groups, providing initial validation of the assumption that STARE can operate on any TACs/brain regions with irreversible kinetics.

STARE also includes an option for vascular correction, where, at the user’s discretion, the source and target TACs can be corrected for any desired level of blood volume fraction. While the level of correlation between STARE K_i_ and corresponding AIF-based estimates is not affected by whether the vascular correction strategy is considered or not within STARE, their agreement slightly varies, and our results suggest that considering the vascular correction strategy within STARE is favorable, at least in this [^18^F]FDG dataset. The suitability of STARE vascular correction strategy should be examined for each tracer and in additional datasets/scanners/populations, and future work could explore optimization of the proposed strategy or implementation of alternative strategies, as well as development of a version of STARE that includes an estimable V_B_ for each brain region.

This is the first presentation of STARE’s theoretical framework and application to PET data; therefore, there are limitations to the conclusions we can draw, as well as many potential future directions of this work. In the current implementation, defining ROIs requires a T1-weighted magnetic resonance imaging (MRI) scan. However, future work with STARE can examine the use of clustering techniques to identify ROIs based only on the PET data. Further, in this first implementation we did not partial volume correct the source and target regions. Future work should explore the effect of partial volume effects on STARE performance, especially in the context of neurodegenerative disorders.

STARE was applied here with six ROIs; however, any region can potentially be included within STARE, with the caveat that, as the number of considered ROIs increase, the free parameter space dimensionality increases (three free parameters per ROI). Further, in our assessment, our proposed strategy of having the source region rotate among the ROI set is favorable with [^18^F]FDG in brain tissue as applied to psychiatric or neurodegenerative disorders, where each region of the brain shows relatively similar kinetics and no source region seems to outperform the other. For other tracers or in other conditions (e.g., [^18^F]FDG uptake in tumor), one region may outperform other regions as a source, or there may be *a priori* reasons for selecting a specific region as the source, in which case, the rotation step can be avoided. Additionally, a weighting scheme could be optimized that weights K_i_ estimates from particular source rotations more heavily, if there is a specific rationale for it, rather than using the unweighted average approach proposed here. Simulated annealing was selected here as the optimizer of choice for STARE. However, simulated annealing is time consuming, taking ~20 minutes for optimization across the six parallelized source rotations used here. Future work might investigate the performance of other optimizers, which may decrease the processing time per scan. A more computationally efficient implementation may also allow STARE to be extended to voxel-level estimation, and involves developing adaptations that are robust in the face of the higher noise level present at the voxel-level. This extension of STARE’s application is part of our future planned work.

In this initial validation, all scans were obtained on the same PET scanner. Validation will be required to assess STARE’s generalizability, as we have implemented it here, to other types of PET scanners and other radiotracers, an important future direction for this work. As shown in the simulations, STARE anchoring is robust to deviations in the signal within the vasculature cluster up to a certain point, but can be hindered by large inaccuracies in the mean signal within the vasculature cluster. For this reason, the two-stage k-means extraction process is critical, because it allows the number of clusters to be automatically selected from the data based on the kinetics and noise within that particular scan, in order to avoid such inaccuracies in the extracted vasculature cluster. However, in the PVC step, the user must supply the FWHM of the point spread function for the specific scanner and reconstruction approach. It is essential that this value is selected appropriately, as grossly altering the AUC of the mean signal in the vasculature cluster, could begin to effect STARE performance.

Although it is not used for this purpose within STARE, this novel, freely available vasculature cluster extraction, can also be implemented independently to potentially generate an IDIF. This will require optimization and validation with external datasets, especially given that the robustness to alterations in the vasculature cluster signal, has been investigated here in the context of total STARE performance, and not in the context of direct effects to kinetic modeling when an IDIF that is subject to effects from scanner FOV and spatial resolution, smoothing during reconstruction, motion, PVC, etc. is used as input. The best future use of the vasculature extraction portion of STARE would likely be in combination with a validated scaling technique. We emphasize that this is an initial validation of STARE as a whole, including both the Source-to-Target tissue model and anchoring portions, and external validation under different scenarios is critical to test STARE’s robustness to other PET scanners, radiotracers, populations, and treatment conditions.

## 5 CONCLUSIONS

STARE - Source-to-Target Automatic Rotating Estimation – is a novel approach for automated, blood-free quantification of the net influx rate (K_i_) of PET radiotracers with irreversible kinetics, that relies solely on the individual’s dynamic PET data. Letting each of the brain regions for which quantification is desired to act, in turn, as a common “source” brain region for all other “target” regions to be expressed as a function of, allows STARE to gain strength by exploiting the information in multiple regions at once with the goal of accurate and precise estimation of K_i_. STARE is “anchored” for participant-specific identifiability using information derived from a novel vasculature cluster extraction and bootstrapping procedure. We validated STARE with a set of [^18^F]FDG scans, for which we showed that K_i_ can be estimated stably, in good agreement with “gold-standard” AIF-based estimates, and with similar performance across diagnostic groups. In simulations, K_i_ estimates were largely robust to characteristics of the vascular cluster of voxels used for STARE anchoring. With more validation, STARE can accelerate use of quantitative PET imaging in the clinic by simplifying acquisition, reducing cost, and affording individualized, non-invasive quantification.

## ** Nonstandard abbreviations

STARE: Source-to-Target Automatic Rotating Estimation

## Declarations of interest

Drs. Bartlett, Zanderigo, and Ogden declare none. Dr. Mann receives royalties from the Research Foundation for Mental Hygiene for commercial use of the C-SSRS.

## Funding

This work was supported by the National Institute of Biomedical Imaging and Bioengineering [R01EB026481, PI: Francesca Zanderigo, PhD].

## Appendix – STARE Derivation

The time activity curves (TACs) in any given source region’s *S*(*C_s_*(*t*)) and any given target region’s *T*(*C_T_*(*t*)) can be expressed as a function of the concentration of radiotracer in arterial plasma (*C_p_*(*t*)) according to the two tissue irreversible (2TCirr) compartment model:

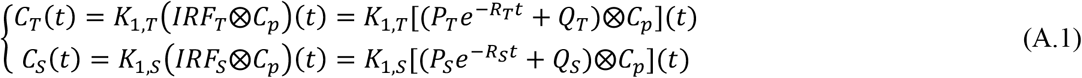

where:

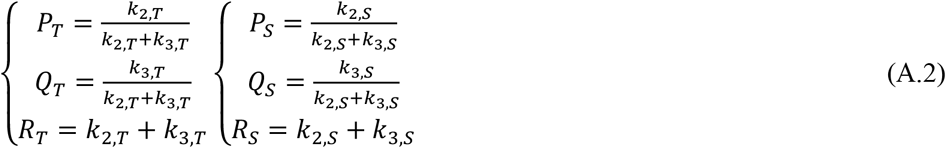

and where *t* is the vector of PET frame time-points, *K*_1,*s*_, *k*_2,*s*_, and *k*_3,*s*_ are the 2TCirr micro-parameters for the source region *S*, *K*_1,*T*_, *k*_2,*T*_, and *k*_3,*T*_ are the 2TCirr micro-parameters for the target region *T*, and *IRF* is the impulse response function in each region.

Transforming the system of equations in (1) into Laplace domain, we obtain:

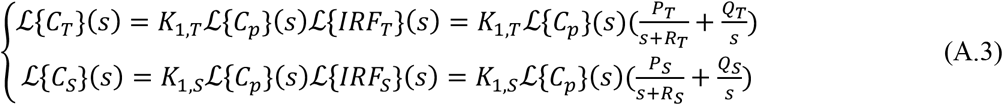

By solving for *C_p_*(*s*) in the second equation in the system in (A.3) and substituting it into the Equation (A.1), we obtain:

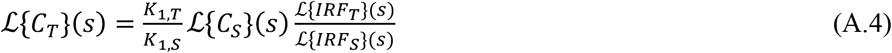

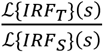 can be expressed as follows:

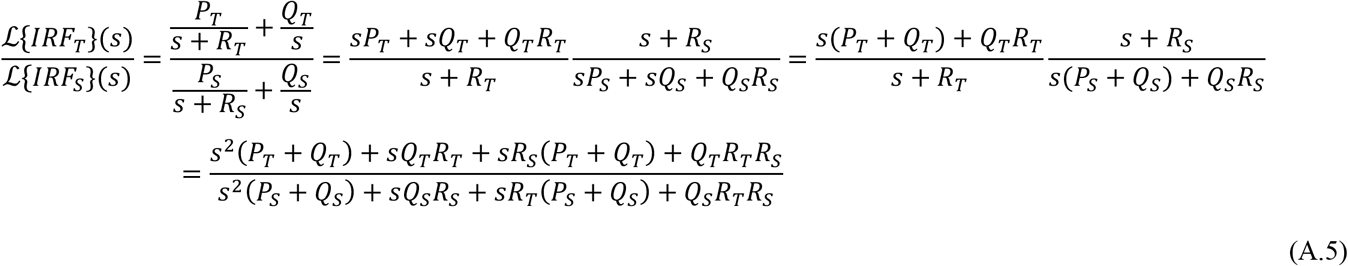

where:

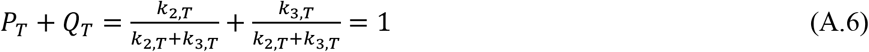

and

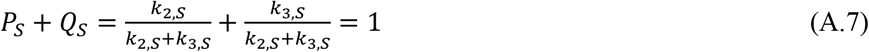

Therefore, simplifying further, this yields:

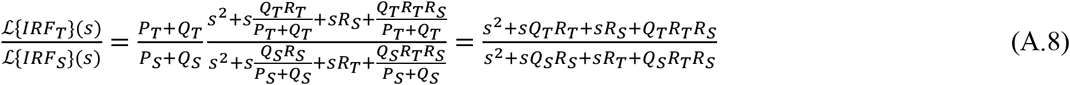

By defining the following system of Equations:

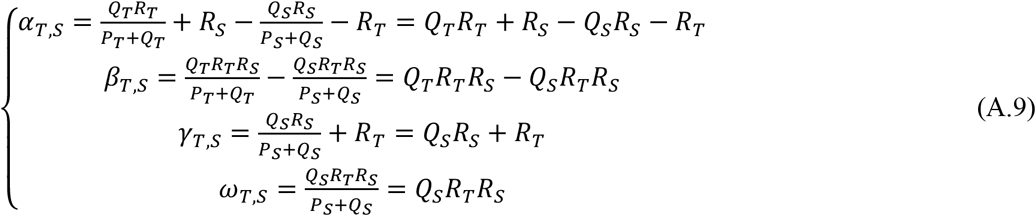

Equation (A.8) can be expressed as:

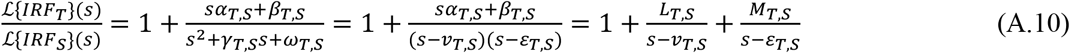

with:

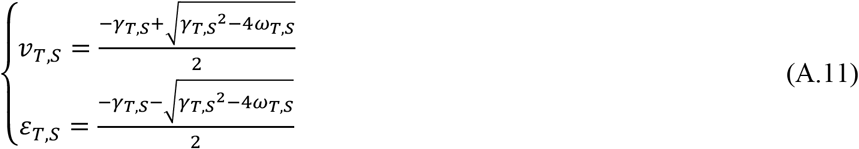

and:

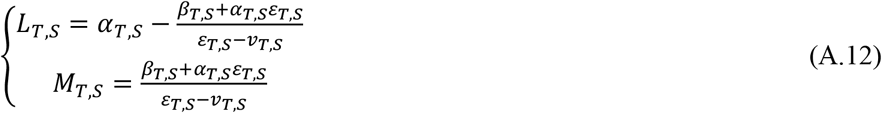

By applying the inverse Laplace transform to Equation (A.10), we obtain a function, *Z_T,s_*(*t*), in the time domain as:

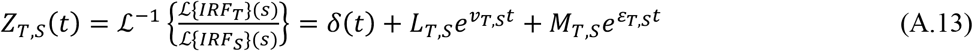

where *δ*(*t*) is the Dirac delta function.

By applying the inverse Laplace transform to Equation (A.4), and considering Equation (A.13), we can express the TAC in each target region (*C_T_*(*t*)) as a function of the TAC in the source region (*C_s_*(*t*)) and of 6 free parameters: 3 for the source region, which are in common across all target regions (*K*_1,*s*_, *k*_2,*s*_, *k*_3,*s*_) and 3 for each target region (*K*_1,*T*_, *k*_2,*T*_, *k*_3,*T*_):

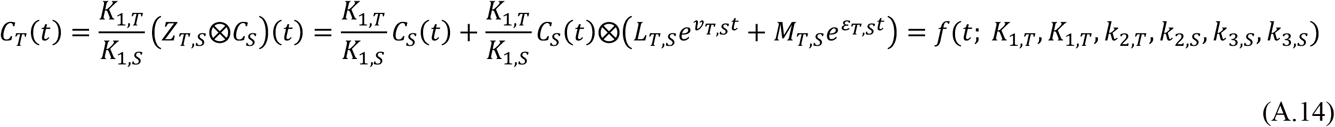

## Notes

### Competing Interest Statement

Drs. Bartlett, Zanderigo, and Ogden declare no competing interests. Dr. Mann receives royalties from the Research Foundation for Mental Hygiene for commercial use of the C-SSRS.

## References

1 Innis, R. B. et al. Consensus nomenclature for in vivo imaging of reversibly binding radioligands. Journal of Cerebral Blood Flow & Metabolism 27, 1533–1539 (2007).

2 Patlak, C. S., Blasberg, R. G. & Fenstermacher, J. D. Graphical evaluation of blood-to-brain transfer constants from multiple-time uptake data. Journal of Cerebral Blood Flow & Metabolism 3, 1–7 (1983).

3 Devanand, D. et al. Pittsburgh compound B (11C-PIB) and fluorodeoxyglucose (18 F-FDG) PET in patients with Alzheimer disease, mild cognitive impairment, and healthy controls. Journal of geriatric psychiatry and neurology 23, 185–198 (2010).

4 Roccia, E. et al. Quantifying Brain [(18)F]FDG Uptake Noninvasively by Combining Medical Health Records and Dynamic PET Imaging Data. IEEE J Biomed Health Inform 23, 2576–2582, doi:10.1109/JBHI.2018.2890459 (2019).

5 Pacák, J., Točík, Z. & Černý, M. Synthesis of 2-deoxy-2-fluoro-D-glucose. Journal of the Chemical Society D: Chemical Communications, 77–77 (1969).

6 Sokoloff, L. et al. The [14C] deoxyglucose method for the measurement of local cerebral glucose utilization: theory, procedure, and normal values in the conscious and anesthetized albino rat 1. Journal of neurochemistry 28, 897–916 (1977).

7 Phelps, M. et al. Tomographic measurement of local cerebral glucose metabolic rate in humans with (F - 18) 2 - fluoro - 2 - deoxy - D - glucose: validation of method. Annals of Neurology: Official Journal of the American Neurological Association and the Child Neurology Society 6, 371–388 (1979).

8 Zasadny, K. R. & Wahl, R. L. Standardized uptake values of normal tissues at PET with 2-[fluorine-18]-fluoro-2-deoxy-D-glucose: variations with body weight and a method for correction. Radiology 189, 847–850 (1993).

9 Boellaard, R. Standards for PET image acquisition and quantitative data analysis. Journal of nuclear medicine (2009).

10 Vriens, D., Visser, E. P., de Geus-Oei, L.-F. & Oyen, W. J. Methodological considerations in quantification of oncological FDG PET studies. European journal of nuclear medicine and molecular imaging 37, 1408–1425 (2010).

11 Westerterp, M. et al. Quantification of FDG PET studies using standardised uptake values in multicentre trials: effects of image reconstruction, resolution and ROI definition parameters. European journal of nuclear medicine and molecular imaging 34, 392–404 (2007).

12 Guo, H., Renaut, R. A. & Chen, K. An input function estimation method for FDG-PET human brain studies. Nucl Med Biol 34, 483–492, doi:10.1016/j.nucmedbio.2007.03.008 (2007).

13 Wong, K.-P., Meikle, S. R., Feng, D. & Fulham, M. J. Estimation of input function and kinetic parameters using simulated annealing: application in a flow model. IEEE Transactions on Nuclear Science 49, 707–713 (2002).

14 Wong, K. P., Feng, D., Meikle, S. R. & Fulham, M. J. Simultaneous estimation of physiological parameters and the input function--in vivo PET data. IEEE Trans Inf Technol Biomed 5, 67–76 (2001).

15 Chen, K. et al. Noninvasive quantification of the cerebral metabolic rate for glucose using positron emission tomography, 18F-fluoro-2-deoxyglucose, the Patlak method, and an image-derived input function. Journal of Cerebral Blood Flow & Metabolism 18, 716–723 (1998).

16 Ogden, R. T., Zanderigo, F., Choy, S., Mann, J. J. & Parsey, R. V. Simultaneous estimation of input functions: an empirical study. Journal of Cerebral Blood Flow & Metabolism 30, 816–826 (2010).

17 Bohorquez, S. S. et al. Flumazenil metabolite measuremente in plasma is not necessary for accurate brain benzodiazepine receptor quantification. Eur. J. Nucl. Med, 1674–1683.

18 Zanotti-Fregonara, P., Chen, K., Liow, J. S., Fujita, M. & Innis, R. B. Image-derived input function for brain PET studies: many challenges and few opportunities. J Cereb Blood Flow Metab 31, 1986–1998, doi:10.1038/jcbfm.2011.107 (2011).

19 Takikawa, S. et al. Noninvasive quantitative fluorodeoxyglucose PET studies with an estimated input function derived from a population-based arterial blood curve. Radiology 188, 131–136, doi:10.1148/radiology.188.1.8511286 (1993).

20 Cunningham, V. J. et al. Compartmental analysis of diprenorphine binding to opiate receptors in the rat in vivo and its comparison with equilibrium data in vitro. Journal of Cerebral Blood Flow & Metabolism 11, 1–9 (1991).

21 Hume, S. P. et al. Quantitation of carbon-11-labeled raclopride in rat striatum using positron emission tomography. Synapse 12, 47–54, doi:10.1002/syn.890120106 (1992).

22 Riabkov, D. Y. & Di Bella, E. V. Estimation of kinetic parameters without input functions: analysis of three methods for multichannel blind identification. IEEE Transactions on biomedical engineering 49, 1318–1327 (2002).

23 Feng, D., Wong, K. P., Wu, C. M. & Siu, W. C. A technique for extracting physiological parameters and the required input function simultaneously from PET image measurements: theory and simulation study. IEEE Trans Inf Technol Biomed 1, 243–254 (1997).

24 Sari, H. et al. Non-invasive kinetic modelling of PET tracers with radiometabolites using a constrained simultaneous estimation method: evaluation with 11 C-SB201745. EJNMMI research 8, 1–12 (2018).

25 Zanderigo, F. et al. [11 C] Harmine Binding to Brain Monoamine Oxidase A: Test-Retest Properties and Noninvasive Quantification. Molecular Imaging and Biology 20, 667–681 (2018).

26 Maroy, R., de Gavriloff, S., Jouvie, C. & Trébossen, R. in IEEE Nuclear Science Symposuim & Medical Imaging Conference. 2081–2083 (IEEE).

27 Phelps, M. E. et al. Tomographic measurement of local cerebral glucose metabolic rate in humans with (F-18)2-fluoro-2-deoxy-D-glucose: validation of method. Ann Neurol 6, 371–388, doi:10.1002/ana.410060502 (1979).

28 Wakita, K. et al. Simplification for measuring input function of FDG PET: investigation of 1-point blood sampling method. J Nucl Med 41, 1484–1490 (2000).

29 Bartlett, E. Less-Invasive and Non-Invasive Quantification of Positron Emission Tomography Data PhD thesis, Stony Brook University, (2019).

30 Takikawa, S. et al. Noninvasive quantitative fluorodeoxyglucose PET studies with an estimated input function derived from a population-based arterial blood curve. Radiology 188, 131–136 (1993).

31 Zhou, Y. et al. Generalized population-based input function estimation given incomplete blood sampling in quantitative dynamic FDG PET studies. Journal of Nuclear Medicine 53, 380–380 (2012).

32 Chen, K. et al. Characterization of the image-derived carotid artery input function using independent component analysis for the quantitation of [18F] fluorodeoxyglucose positron emission tomography images. Physics in Medicine & Biology 52, 7055 (2007).

33 Naganawa, M. et al. Extraction of a plasma time-activity curve from dynamic brain PET images based on independent component analysis. IEEE transactions on biomedical engineering 52, 201–210 (2005).

34 Zanotti-Fregonara, P. et al. Comparison of eight methods for the estimation of the image-derived input function in dynamic [18F]-FDG PET human brain studies. Journal of Cerebral Blood Flow & Metabolism 29, 1825–1835 (2009).

35 Naganawa, M. et al. Assessment of population-based input functions for Patlak imaging of whole body dynamic 18 F-FDG PET. EJNMMI physics 7, 1–15 (2020).

36 Wang, B., Ruan, D. & Liu, H. Noninvasive Estimation of Macro-parameters by Deep learning. IEEE Transactions on Radiation and Plasma Medical Sciences (2020).

37 Bartlett, E. A. et al. Quantification of Positron Emission Tomography Data Using Simultaneous Estimation of the Input Function: Validation with Venous Blood and Replication of Clinical Studies. Mol Imaging Biol, doi:10.1007/s11307-018-1300-1 (2018).

38 Wakita, K., Imahori, Y., Ido, T. & Fujii, R. Simplification for measuring input function of FDG PET: investigation of 1-point blood sampling method. The Journal of Nuclear Medicine 41, 1484 (2000).

39 Chiou, W. L. The phenomenon and rationale of marked dependence of drug concentration on blood sampling site. Clinical pharmacokinetics 17, 175–199 (1989).

40 Sari, H. et al. Estimation of an image derived input function with MR-defined carotid arteries in FDG-PET human studies using a novel partial volume correction method. J Cereb Blood Flow Metab 37, 1398–1409, doi:10.1177/0271678X16656197 (2017).

41 Zanotti-Fregonara, P., Chen, K., Liow, J.-S., Fujita, M. & Innis, R. B. Image-derived input function for brain PET studies: many challenges and few opportunities. Journal of Cerebral Blood Flow & Metabolism 31, 1986–1998 (2011).

42 Pincus, M. Letter to the editor—a Monte Carlo method for the approximate solution of certain types of constrained optimization problems. Operations research 18, 1225–1228 (1970).

43 Leenders, K. et al. Cerebral blood flow, blood volume and oxygen utilization: normal values and effect of age. Brain 113, 27–47 (1990).

44 Devanand, D. P. et al. Pittsburgh compound B (11C-PIB) and fluorodeoxyglucose (18 F-FDG) PET in patients with Alzheimer disease, mild cognitive impairment, and healthy controls. J Geriatr Psychiatry Neurol 23, 185–198, doi:10.1177/0891988710363715 (2010).

45 Graham, M. M. Physiologic smoothing of blood time-activity curves for PET data analysis. Journal of Nuclear Medicine 38, 1161–1168 (1997).

46 Bates, D., Mächler, M., Bolker, B. & Walker, S. Fitting linear mixed-effects models using lme4. arXiv preprint arXiv:1406.5823 (2014).

47 Team, R. C. (Vienna, Austria, 2013).

